# Cryptochrome 4b protein is likely irrelevant for the radical pair based magnetoreception in the European robin

**DOI:** 10.1101/2025.02.21.639466

**Authors:** Jingjing Xu, Alisha Bhanu Pattani Ameerjan, Jonathan Hungerland, Georg Langebrake, Tina Ravnsborg, Ole N. Jensen, Jessica Schmidt, Rabea Bartölke, Takaoki Kasahara, Baladev Satish, Leonard Schwigon, Karin Dedek, Arne Nolte, Miriam Liedvogel, Ilia A. Solov’yov, Henrik Mouritsen

**Affiliations:** Department of Biochemistry and Molecular Biology, University of Southern Denmark, Campusvej 55, 5230 Odense M, Denmark; Institute of Biology and Environmental Sciences, University of Oldenburg, Carl-von-Ossietzky-Str. 9-11, 26129 Oldenburg, Germany; Institute of Physics, University of Oldenburg, Carl-von-Ossietzky-Str. 9-11, 26129 Oldenburg, Germany; Institute of Avian Research, An der Vogelwarte 21, 26386 Wilhelmshaven, Germany; Center for Nanoscale Dynamics (CENAD), University of Oldenburg, Ammerländer Heerstr. 114-118, 26129 Oldenburg, Germany; Research Centre for Neurosensory Science, University of Oldenburg, Carl-von-Ossietzky-Str. 9-11, 26129 Oldenburg, Germany

## Abstract

Avian cryptochrome 4 (Cry4) protein is a putative magnetosensitive molecule facilitating precise long-distance navigation in migratory birds. Two splice variants of Cry4 were reported in European robin (*Erithacus rubecula*), namely *Er*Cry4a and *Er*Cry4b. It is known that *Er*Cry4a protein exhibits electron transfer between the flavin adenine dinucleotide (FAD) cofactor and tryptophan residues that generates magnetically sensitive radical pairs for magnetoreception. However, little is known about the *Er*Cry4b isoform. We therefore characterized the properties of *Er*Cry4b to see whether it fulfills prerequisites to be a radical pair magnetic sensor molecule. Our results show that *Er*Cry4b protein does not bind FAD *in vitro*. Computational structure simulations revealed that the FAD non-binding in *Er*Cry4b is likely due to protein structure dynamics. Furthermore, *Er*Cry4b protein abundance in the robin retina, cerebellum and liver is below the detection limit of immunoprecipitation assays coupled with mass spectrometry. Meanwhile, transcript analyses show that *ErCRY4b* mRNA abundance is 10 times less than *ErCRY4b* in the retina. In conclusion, *Er*Cry4b does not fulfill the prerequisites to be a radical pair based magnetic sensing molecule due to the lack of FAD binding, and it might not even be expressed as a functional protein in the European robin.

## Introduction

The phenomenon of night-migratory birds navigating long distances with remarkable precision has captivated scientists for decades. One of the prominent navigation cues in nature is the Earth’s magnetic field. Migratory birds can use the Earth’s magnetic field as a compass for directional guidance, and as a map for a precise positional reference [1, 2, 3, 4, 5, 6, 7, 8]. How birds sense magnetic fields remains an intriguing scientific question in sensory biology [9]. Over the last decades, accumulating theoretical and experimental evidence has supported a quantum light-induced radical pair mechanism in the avian retina [10, 11, 12, 13, 14, 15, 16]. The radical pair mechanism of magnetoreception requires a light sensitive protein which can generate radical pairs upon short-wavelength light excitation [17]. Cryptochrome (Cry) has been proposed to be such a magnetosensory protein as it is the only currently known animal photopigment that can produce long-lived and spin-correlated radical pairs [12, 17, 18, 19, 20].

*Cryptochromes (CRYs)* are a multigene family of blue light photoreceptors, mostly discussed in the context of circadian rhythm regulation. So far, three different genes have been reported in birds and each has two splice variants: *CRY1a, CRY*1b, *CRY*2a, *CRY*2b, *CRY*4a and *CRY*4b [19, 21, 22, 23, 24, 25, 26, 27, 28, 29]. Among the six isoforms, *CRY*4a showed a seasonal mRNA expression pattern: European robins (*Erithacus rubecula*) *CRY*4a (*ErCRY4a*) was expressed approximately 2.5 times higher during the migratory seasons than that during non-migratory seasons in the retina [24]. In comparison, the other *CRY* isoforms show diurnal circadian oscillation patterns [24, 25, 28]. Moreover, *Er*Cry4a protein is expressed in double cone photoreceptor cells in the retina, which shows a very pronounced 180° mosaic in the periphery of the retina and a less pronounced 90°mosaic in the central retina [24, 30, 31]. The peripheral 180° orientation of neighboring double cones could facilitate to the extraction of directional information derived from the Earth’s magnetic field independent of light intensity and polarization angle variation [32].

The core of the radical pair mechanism of magnetoreception is the flavin adenine dinucleotide (FAD) chromophore in a cryptochrome protein [17, 33], because FAD initiates electron transfer and forms radical pairs with the protein. Several independent studies have shown that avian Cry4a proteins bind FAD as a cofactor [34, 35, 36, 37]. Specifically, *Er*Cry4a exhibits electron transfer from tryptophan (Trp) residues to the FAD cofactor, forming spin-correlated [FAD^*•−*^ TrpH^*•*+^] radical pairs, which are sensitive to external magnetic fields [37, 38, 39]. Therefore, Cry4a protein is considered the most likely magnetosensory molecule in birds.

While *Er*Cry4a has been characterized as magnetosensory molecule, it is unknown whether the alternative splice variant *Er*Cry4b could play any role in magnetoreception. *Er*Cry4b amino acid sequence is identical to *Er*Cry4a except for one additional exon, which locates in the 2^nd^ intron of *ErCRY4a* gene (figure 1a) [40]. *ErCRY4b* mRNA is considered to be caused by alternative splicing. Alternative splicing in general is a gene expression regulation mechanism that allows a single gene to produce multiple mRNA isoforms, thereby greatly increasing proteome complexity [41, 42]. On one hand, *Er*Cry4b protein sequence maintains an intact FAD binding domain, which could in principle facilitate FAD binding and form radical pairs. On the other hand, the latest avian genome analyses showed that the *CRY4b*-specific exon carries loss-of-function mutations (e.g., stop codons), a pattern characteristic for intronic regions [29]. This poses the question whether the *ErCRY*4b mRNA isoform is translated into a functional protein product *in vivo*.

**Figure 1.**
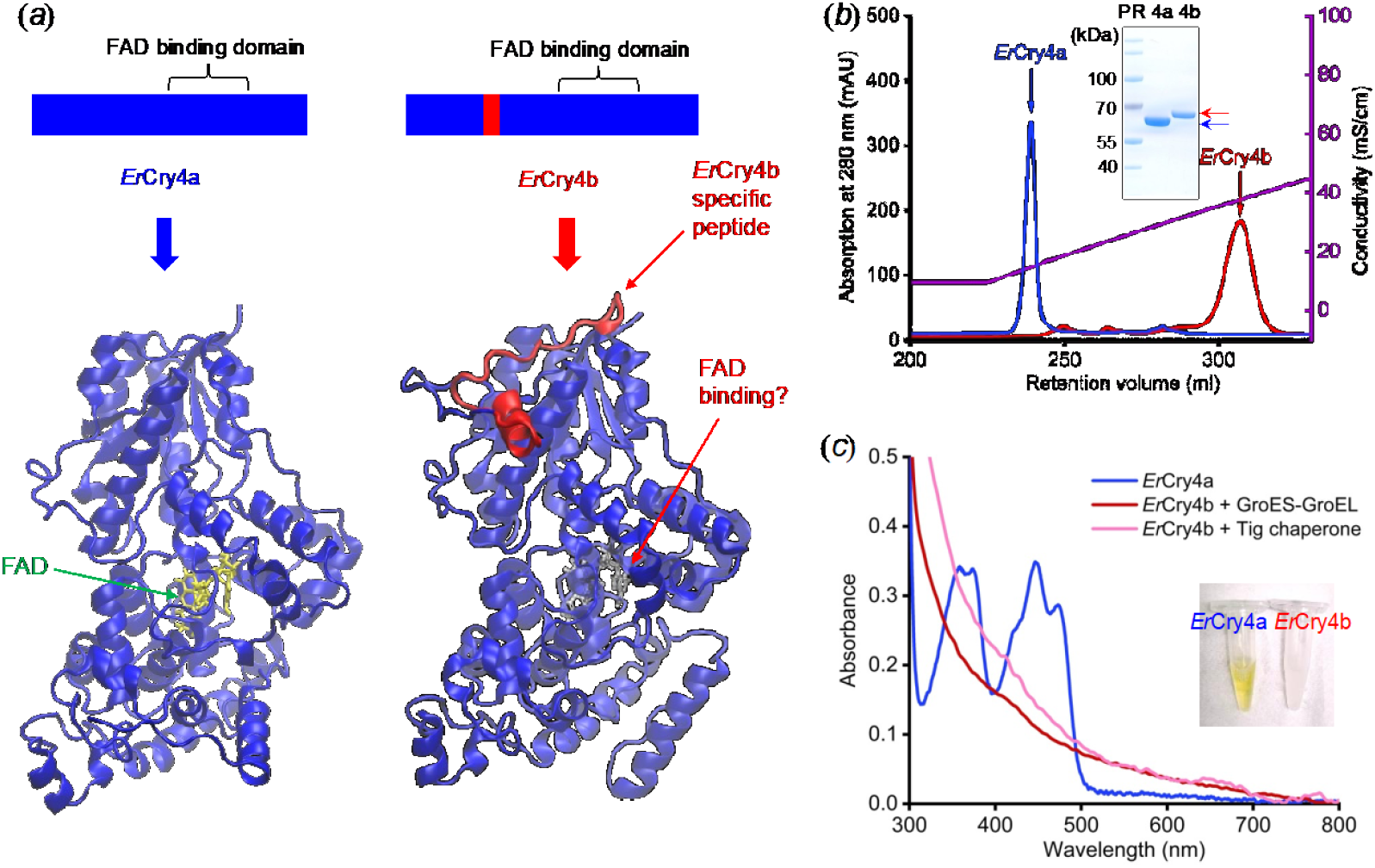
European robin (*Erithacus rubecula*) Cryptochrome 4b protein does not bind FAD *in vitro*. (*a*) Illustration of 2D and 3D structures of European robin Cryptochrome 4a and 4b proteins (*Er*Cry4a and *Er*Cry4b). Both proteins contain the conserved flavin adenine dinucleotide (FAD) binding domain and *Er*Cry4a is known to bind FAD. *Er*Cry4a protein and its conserved amino acid are marked in blue, while *Er*Cry4b unique peptide is highlighted in red. The color scheme introduced here is maintained throughout the article: blue and red represents *Er*Cry4a and *Er*Cry4b, respectively. The question mark indicates the primary focus of the present study: the FAD binding status in *Er*Cry4b. (*b*) Anion exchange chromatography profiles of *Er*Cry4a and *Er*Cry4b. The solvent conductivity along the retention volume is shown in the purple color. While *Er*Cry4a was eluted at a low ionic strength of ca. 20 mS/cm, *Er*Cry4b was eluted at a high ionic strength corresponding to the conductivity of ca. 37 mS/cm. The inset is an Coomassie blue staining image of purified *Er*Cry4a and *Er*Cry4b on a sodium dodecyl sulfate–polyacrylamide gel, showcasing the molecular weight difference between the two proteins. (*c*) UV-visible absorption spectrum of *Er*Cry4a and *Er*Cry4b. *Er*Cry4a served as a reference. The absorbance peak at 450 nm with shoulders is the characteristic of FAD binding in a cryptochrome protein (blue). *Er*Cry4b protein did not show such clear peaks despite of recombinant expressions with GroES-GroEL chaperone (red) or Tig chaperone (pink). The inset is a photo of *Er*Cry4a and *Er*Cry4b protein samples. Compared to the yellow sample of *Er*Cry4a, the colorless *Er*Cry4b sample indicates no FAD binding.

To assess whether *Er*Cry4b could be relevant to the radical pair mechanism of magnetoreception, we characterize *Er*Cry4b by combining biochemical assays with protein dynamics simulations and transcript analysis. Specifically, the objective of the present study is characterizing the FAD binding status *in vitro*, estimating the FAD binding ability by simulating the protein dynamics, detecting *Er*Cry4b protein in bird tissue and screening *CRY4b* transcript in a robin transcriptome database.

## Results

### Non-FAD binding in *Er*Cry4b protein *in vitro*

In the context of radical pair based magnetoreception, FAD binding is necessary for generating flavin radicals and thus to allow for putative magnetic sensing in a cryptochrome protein [17]. Thus, our primary focus here was to examine whether *Er*Cry4b protein binds FAD. Towards this goal, we expressed and purified *Er*Cry4b protein using bacterial recombinant protein expression methods upon modification on the previous protocol [37]. The results showed that *Er*Cry4b was not as soluble as *Er*Cry4a. The majority of *Er*Cry4b protein fell into insoluble fraction after expression (figure S1-S2). Soluble *Er*Cry4b protein was only obtained with the aid of GroES-GroEL chaperone co-expression. Upon the chaperone optimization, *Er*Cry4b protein expression yield was up to ca. 3 mg per liter *E. coli* cell culture, which was comparable to that for *Er*Cry4a [37]. The purified *Er*Cry4b protein (66 kDa) showed a clear mass shift compared to *Er*Cry4a (63 kDa) on SDS polyacrylamide gel (figure 1*b*). Notably, *Er*Cry4b protein was eluted by a higher solvent conductivity (ca. 37 mS/cm) compared to *Er*Cry4a (ca. 20 mS/cm) in an anion exchange chromatography (figure 1*a*). The different elution profile of *Er*Cry4b compared to that of *Er*Cry4a suggests that *Er*Cry4b may carry a different surface charge pattern due to a different protein conformation.

The characteristic feature of FAD binding is an absorption peak at 450 nm with surrounding small fine peak structures [37, 43, 44, 45]. If *Er*Cry4b binds FAD, *Er*Cry4b UV-visible light absorption spectrum should show the FAD binding characteristic feature comparable to that in *Er*Cry4a. However, such characteristic was absent from *Er*Cry4b UV-visible absorption spectrum (figure 1*c*). Instead, *Er*Cry4b UV-visible spectra showed a slope-like profile across the wavelength range from 300 nm to 800 nm, indicating no FAD binding. Moreover, the slope-like profile may be attributed to baseline artifacts from protein aggregation [46]. The *Er*Cry4b protein aggregation phenomenon indicated a less stable structure of *Er*Cry4b compared to *Er*Cry4a in the same pH8 solvent buffer. Most importantly, the purified *Er*Cry4b protein sample was colorless, further supporting the absence of bound FAD in eluted *Er*Cry4b (see the inset of figure 1*c*). In combination, the biochemistry results indicated that the recombinant expressed *Er*Cry4b protein did not bind FAD *in vitro* or that it was not well folded *in vitro*, which might in turn hamper FAD binding.

### The dynamics of *E*rCry4b FAD binding domain

To examine FAD binding affinity in *Er*Cry4b protein, we have simulated the interaction energy between FAD and *Er*Cry4b protein scaffold, as well as the overall conservation of the protein structure during its dynamical evolution. Figure 2*a* shows the probability distribution of the total sum of the total electrostatic and van-der-Waals interaction energy arising between the FAD cofactor and the protein or the surrounding solvent. The average potential energy of FAD in *Er*Cry4b is less favorable by -39 kcal/mol compared to that of FAD in *Er*Cry4a (solid lines in figure 2*a*). Notably, FAD in the pocket of *Er*Cry4b has stronger interactions with the solvent than that in *Er*Cry4a (dash lines in figure 2*a*). The high solvent access indicates an increased binding energy barrier for FAD in *Er*Cry4b.

**Figure 2.**
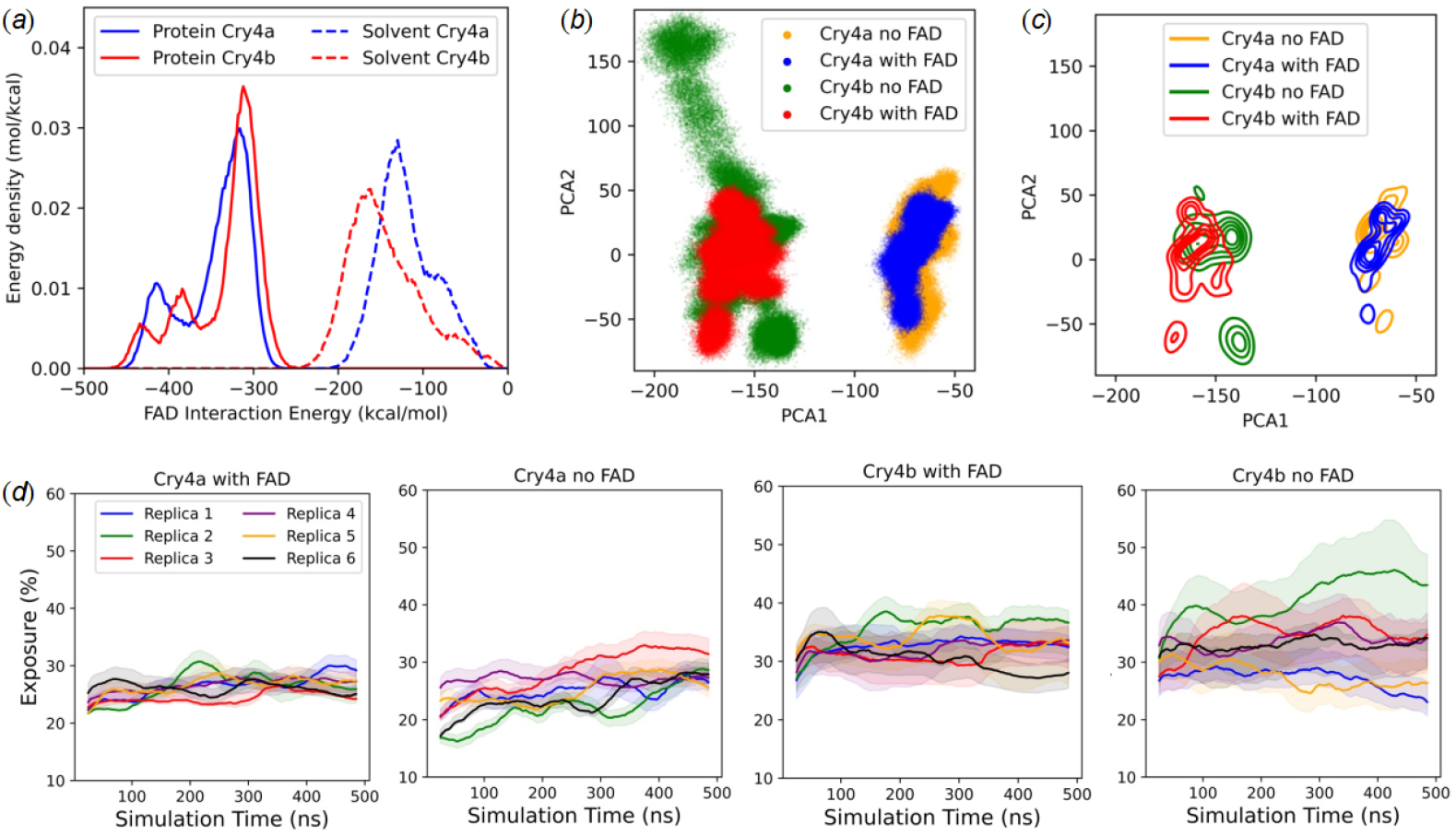
Computational simulations suggest distinct differences in FAD binding energy, protein trajectories, and solvent accessibility of the FAD binding pocket between *Er*Cry4a and *Er*Cry4b. (*a*) Probability distribution of the FAD interaction energy with the protein scaffold (solid lines) and the solvent (dashed lines). (*b*) Principal component analysis (PCA) of *Er*Cry4a and *Er*Cry4b protein conformations with/without FAD. Each dot represents scattered datapoints along the first two components of the resulting PCA in the chosen coordinate intersection. (*c*) Contour plots of the density with equi-spaced lines in the PCA space. The top-left data points in (b) are not represented in (*c*) due to highly sparsity. (*d*) Time-evolution of the solvent exposure of residues that are found within a solvent shell around the FAD after structure minimization for *Er*Cry4a and *Er*Cry4b with/without FAD, respectively. The data was averaged using a sliding window of ±25 ns. The line shows the median value over different solvent shell cutoffs, while the color shades show the range of exposure that is obtained from different solvent shell cutoffs. Each color is a different replica simulation of the respective proteins.

Further investigation of the protein dynamics supports the FAD binding affinity difference between *Er*Cry4a and *Er*Cry4b. One of the essential requirements for FAD binding is the dynamically preserved existence of the FAD binding pocket. To illustrate consistency in the dynamic evolution of *Er*Cry4a protein independent of FAD presence, we carried out a principal component analysis (PCA) (see SI for details). The PCA results show that *Er*Cry4a apoprotein and holoprotein conformations (orange: apoprotein without FAD; blue: holoprotein with FAD) overlap in the space spanned in the principal component analysis (PCA) (figure 2*b*). The dominant local density maxima of *Er*Cry4a apoprotein without FAD appear to be transition states between the density maxima of *Er*Cry4a with FAD (figure 2*c*). In contrast, *Er*Cry4b apoprotein and holoprotein conformations (orange: apoprotein without FAD; red: holoprotein with FAD) differ with only little overlap and explore more distant regions of the principal component space (figure 2*b*). The contour plot also reveals the correspondingly separate density maxima for *Er*Cry4b apoprotein and holoprotein (figure 2*c*). For an absolute comparison of the differences between the trajectories, a Wasserstein distance analysis was performed (see figure S6 and Table. S2). The trajectories of *Er*Cry4a with and without FAD showed a distance of 1.51 Å /atom and the trajectories of *Er*Cry4b with and without FAD showed a distance of 1.78 Å/atom (Table. S2). The results indicate that *Er*Cry4a structure is dynamically preserved independent of the presence of FAD, while *Er*Cry4b structure changes depending on the presence of FAD. The root-mean-square deviation (RMSD) analysis suggests that the additional amino acid residues in *Er*Cry4b causes an additional flexible region compared to *Er*Cry4a (figure S7 and S8). *Er*Cry4b does not show a dynamic conservation of its structure independent of the presence of FAD, *Er*Cry4b is more problematic to fulfill the essential requirement of FAD binding.

Another more local measurement of the properties of the FAD binding pocket is the relative solvent exposure of the residues around FAD. In *Er*Cry4a, the relative solvent exposure is rather low around 25% - 30%. In contrast, *Er*Cry4b has a higher relative solvent exposure of 30% - 40% in the presence of FAD and even as high as 20% - 55% in the absence of FAD (figure 2*d*). The high solvent exposure of *Er*Cry4b FAD binding pocket occurs in the phosphate binding loop and the C-terminal region, both of which are expected to be rather flexible (Schuhmann et al., 2024). The results on solvent exposure further indicate towards a comparatively unstable FAD binding pocket for *Er*Cry4b.

### The *Er*Cry4b unique peptide is not detectable *in vivo*

As *Er*Cry4b protein does not bind FAD *in vitro*, we moved forward to investigate Cry4b protein *in vivo*. An antibody was used to enrich *Er*Cry4 proteins from robin tissue lysates. The antibody was validated to be able to recognize both *Er*Cry4a and *Er*Cry4b (figure S2). The Western blot analysis of immunoprecipitation samples from European robin retina shows one single Cry4 band (figure S3). The molecular weight of the protein band is close to 55 kDa, very likely reflecting Cry4a protein.

To confirm whether the protein band corresponded to *Er*Cry4a or *Er*Cry4b, we identified the peptides in the tissue immunoprecipitation samples using mass spectrometry (MS). Recombinantly expressed and purified *Er*Cry4b protein was used as a reference. We first analyzed common peptides shared by *Er*Cry4a and *Er*Cry4b. For example, one of the common peptides is NLTAEDFQR. Theoretical mass over charge ratio (m/*z*) of the doubly charged common peptide is 547.2673. We observed ions of similar m/*z* and in purified recombinant *Er*Cry4b protein sample (547.2687) and the retina immunoprecipitation sample (547.2685). The ions were assigned to the common peptide NLTAEDFQR by Mascot search using Proteome Discoverer.

Second, we looked for *Er*Cry4b unique peptide VKDVLCLALHEER. The cysteine amino acid was modified with a carbamidomethyl group, and one missed cleavage occurred. Under this circumstance, the theoretical m/*z* of a doubly charged VKDVLCLALHEER is 791.4244. In the recombinantly expressed *Er*Cry4b protein sample, the *Er*Cry4b unique peptide was eluted at 37.04 minutes with a m/*z* of 791.4243, which is close to the theoretical m/*z* value (figure 3*c*). The confidence of peptide spectra match was high. 100% *y* ions and 58% *b* ions were matched to theoretical calculations. Thus, the peptide was established as an indicator for the presence of *Er*Cry4b protein.

**Figure 3.**
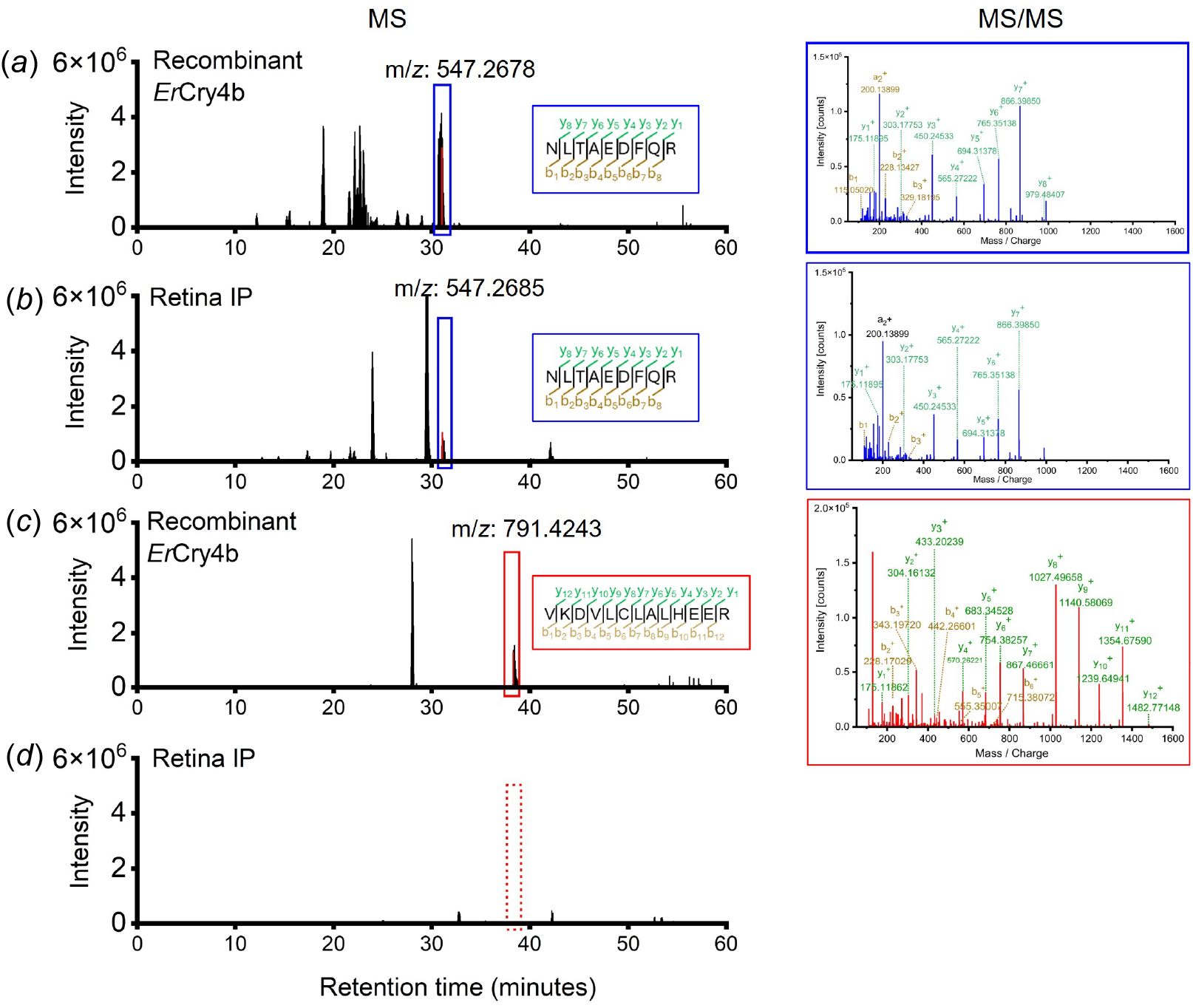
*Er*Cry4b unique peptide was not detectable in robin tissue immunoprecipitation samples. The blue and red solid-line rectangles mark the ions for the common peptide and *Er*Cry4b unique peptide, respectively. Tandem mass spectrometry (MS/MS) data was shown on the right, showcasing the fragment match spectrum of the detected peptides in each sample. (*a*) and (*b*) The common peptide of *Er*Cry4a and 4b (NLTAEDFQR) was detected in a purified *Er*Cry4b protein sample (*a*), and robin retina immunoprecipitation samples (*b*). Theoretical mass over charge ratio (m/*z*) of the doubly charged peptide is 547.2673. Measured m/*z* ratios are 547.2687 and 547.2685 in the purified *Er*Cry4b sample and retina immunoprecipitation sample, respectively. The m/*z* range of the figure is 547.26 - 547.27. (*c*) and (*d*) *Er*Cry4b unique peptide (VKDVLCLALHEER) was clearly detected in a purified *Er*Cry4b protein sample (*c*), but not in immunoprecipitation samples from robin retina (*d*). Theoretical mass over charge ratio (m/*z*) of the doubly charged peptide is 791.4244. The m/*z* range of the figure was 791.40 - 791.44.

Third, we examined the MS spectra of tissue immunoprecipitation samples. If robin tissues express *Er*Cry4b protein at a substantial level, an MS signal close to 791.4243 should be observed at ca. 37.04 minutes in the MS spectra of the robin tissue immunoprecipitation sample. However, such signal was absent in any of the tissue immunoprecipitation samples (figure 3*d* and figure S5).

The presence of the *Er*Cry4a & 4b common peptide and the absence of the *Er*Cry4b unique peptide suggests that *Er*Cry4a is probably the only protein product of *CRY4* gene in retina, cerebellum and liver. The results indicate that *Er*Cry4b protein is either not expressed as abundantly as *Er*Cry4a or not *in vivo* expressed in the three tissues at all.

### Low abundance of *Er*Cry4b mRNA transcript in different tissues

Consistent with the protein mass spectrometry results, PacBio Iso-Seq analyses further revealed a low abundance of *ErCRY4b* mRNA. In the retina, *ErCRY4b* mRNA was 10 times less abundant than *ErCRY4b* mRNA (absolute numbers: 1 and 12, respectively, see figure 4). In addition, we identified six new variants (n1 - n6) with an intron retention or alternative acceptor/donor sites. However, stop codons appears in the retained introns of n1 - n6 variants. Thus, these variants are impossible to be translated into proteins. These transcripts can ultimately be either subject to delayed splicing or are discarded through nuclear degradation or nonsense mediated decay [47]. The most diverse *ErCRY4* splice variants and intron retention variants were found in skin.

**Figure 4.**
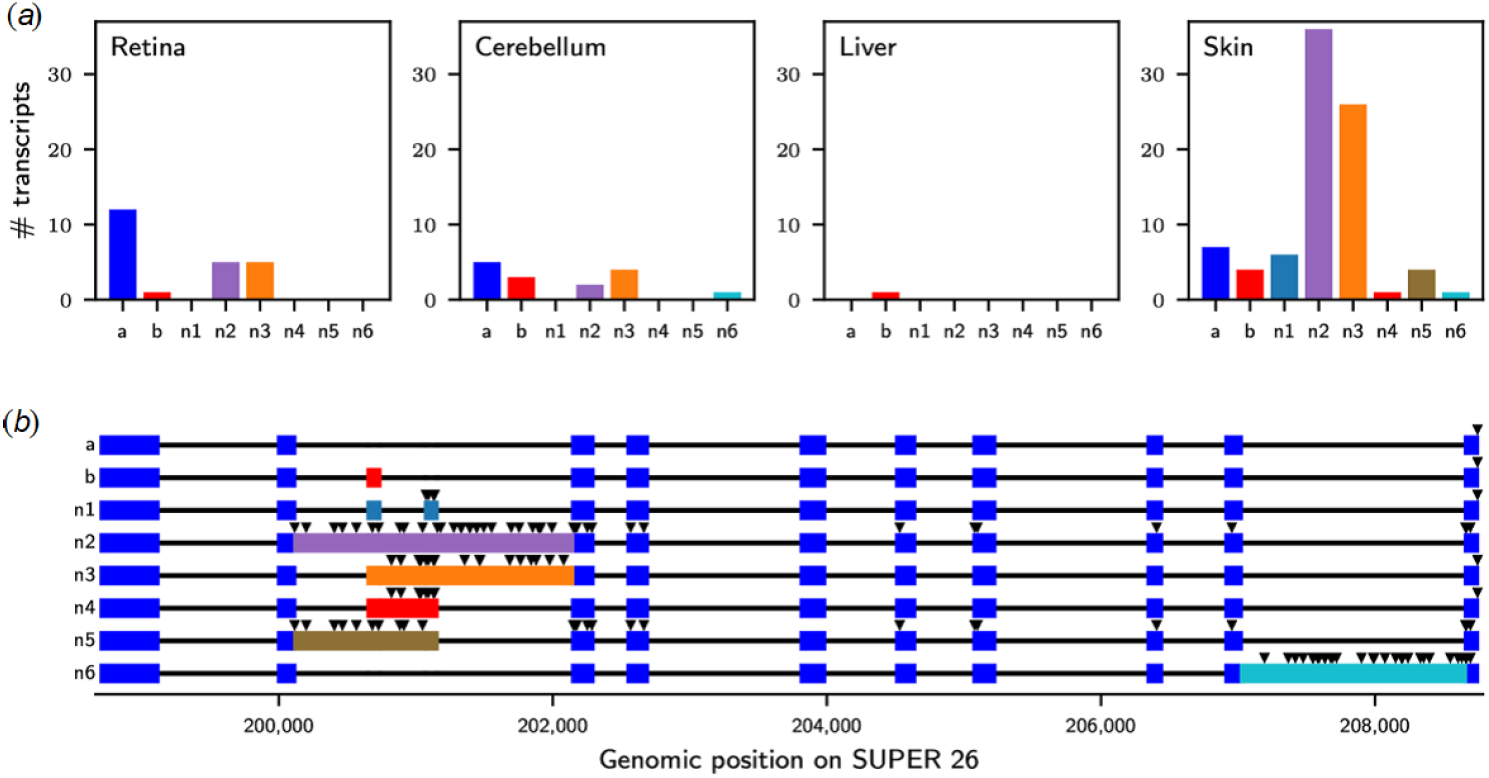
The diverse *ErCRY4* mRNA splice variants in European robin tissues. (*a*) Abundance of *ErCRY4* splice variants (a and b) and newly identified intron retention variants (n1 - n6) in different tissues of European robins. (*b*)Visualization of *ErCRY4* genomic sequence with highlighted exons in colored boxes and intronic regions indicated as black line. The black triangles refer to stop codons. Most of the newly identified variants have intronic retention in the region of the *ErCRY4b* exon.

The abundance ratio of *ErCRY4a* to other variants was used to estimate *ErCRY4* gene splicing efficiency in different tissues. The ratio was 1.09 in retina, compared to 0.42 in brain, 0.86 in lung, 0.08 in skin, 0.6 in testis and 0.4 in heart (see supplementary material for full table of transcript types and numbers). In muscle and liver, only a single transcript of *ErCRY4* was detected, which was too low to calculate a ratio. The lowest ratio was found in skin, which was also where the highest diversity of alternative splicing products was found, suggesting relaxed splicing of *ErCRY4* in the skin.

## Discussion

The present study revealed distinct properties of *Er*Cry4b compared to *Er*Cry4a *in vitro*. The direct observation was that *Er*Cry4b did not bind FAD in *vitro*. FAD binding is one of the essential prerequisites for being a radical pair based magnetic sensing molecule [17, 48]. Without FAD binding, there can be no light-absorption, no electron transfer, no radical pairs and thus no magnetic sensitivity. Following the performed analysis, *Er*Cry4b is unlikely to fulfill the prerequisites being a magnetic sensing molecule.

The non-FAD binding in *Er*Cry4b can be attributed to protein conformation dynamics. If *Er*Cry4b shares a similar folding pattern as *Er*Cry4a, *Er*Cry4b should have similar isoelectric points. In a pH 8 solvent, both *Er*Cry4a and *Er*Cry4b proteins should carry similar amount of net negative charge of around 6.5, bind to a positively charged anion exchange resin with similar affinities, and be eluted by a solvent of similar ionic strength/conductivity. However, our results showed that *Er*Cry4b was unexpectedly eluted at a higher conductivity (37 mS/cm) compared to *Er*Cry4a (20 mS/cm). It indicates that *Er*Cry4b folded differently compared to *Er*Cry4a. The results challenge the assumption that *Er*Cry4b adopts the same conformation as *Er*Cry4a *in vitro*.

Further computational analyses suggest that even if *Er*Cry4b was folded similarly to *Er*Cry4a, its ability to bind FAD is profoundly smaller than that of *Er*Cry4a. The evidence accumulates from interaction energy studies, modelling of protein dynamics and solvent exposure simulation. The binding pocket for FAD in *Er*Cry4b turned out to be energetically less favorable than in the case of *Er*Cry4a. *Er*Cry4b protein dynamics is different once FAD is present or absent. The FAD cofactor in *Er*Cry4b tends to interact more frequently with the solvent than that in *Er*Cry4a, likely indicating an increased binding energy barrier in *Er*Cry4b. Therefore, even if FAD was found in *Er*Cry4b, its retention time is expected to be orders of magnitude smaller than in the case of *Er*Cry4a.

Moreover, *Er*Cry4b protein was not detectable in the European robin retina, despite using the protein enrichment technique of immunoprecipitation and the peptide detection technique of high-resolution mass spectrometry methods. This implies that *Er*Cry4b is expressed at extremely low abundance levels or may not be expressed as a protein product in the retina at all. We do not yet know if *Er*Cry4b or other alternative splicing variants are completely meaningless in other tissues. Therefore, we cannot completely exclude *Er*Cry4b protein expression and function. However, the absence of detectable *Er*Cry4b in the retina contrasts with expectations for a functional sensory protein, which would typically be produced at sufficient levels to perform its biological role.

It is not uncommon that splicing variants do not produce proteins [49]. While transcript data showed that more than 90% of human genes undergo alternative splicing, proteomics studies strongly suggest that most genes have a single dominant splice isoform as a protein [50, 51, 52]. For example, human *CRY1* gene (Ensemble ID: ENSG00000008405) has 5 different transcripts, but only one protein product.

Our transcript analysis suggests that *ErCRY4b* mRNA abundance is 10 times lower than *ErCRY4a*. Previously, Einwich and colleagues reported that *ErCRY4b* exhibited a comparable mRNA expression level as *ErCRY4a* in the robin retina [25]. The seemingly contradictory results may at least partly be explained by different methodology used for mRNA characterization. Einwich and colleagues detected mRNA expression level using quantitative-PCR (qPCR), which relied on the DNA primer specificity to the targeted sequence. The DNA primers used in the previous study [40] recognized not only Cry4b transcripts but also many other splice variants. In total, the *CRY4b* primers from Einwich et al. (2020) should allow to amplify nine *ErCRY4* alternative transcripts from robin retina transcriptome. Summing up the abundance of multiple transcripts can lead to the same order of magnitude in total expression as *ErCRY4* abundance. Thus, the qPCR results in Einwich *et al*. (2020) study showed a seemingly comparable mRNA expression level for *ErCRY4*a and *ErCRY4b*.

In this light, the diurnal rhythm expression pattern of ‘*ErCRY4b*’ essentially represents the expression of multiple mRNA transcripts. Thus, Einwich et al. (2020) data suggests a diurnal rhythm of the efficiency of *CRY4* gene splicing. The *CRY4* gene expression is potentially regulated by different gene-splicing efficiency from day to night. During the night, less non-functional transcripts with introns present are produced as a form of expression regulation. Our transcript analysis also suggests that the splicing efficiency is highest in the retina compared to the other sampled tissues, with a ratio of fully spliced to un-spliced transcripts greater than one. This further corroborates the importance of *ErCRY4a* in the retina and strengthens the circumstantial evidence for its role in magnetoreception.

Notably, our current transcript analysis aligns with the previous phylogenomic study characterizing different *CRY* gene across the avian clade, which revealed that many bird species carry stop codons within the *CRY4b* exon [29]. Together, we suggest that *ErCRY4b* has noncoding roles in at least some lineages and thus may be functionally irrelevant to magnetoreception.

## Conclusion

In summary, *Er*Cry4b protein does not bind the essential cofactor FAD *in vitro* and is not expressed in robin retina. Therefore, *Er*Cry4b is likely irrelevant to the radical pair based magnetoreception. *Er*Cry4a remains the most likely radical pair based magnetic sensory molecule in European robins.

## Methods and materials

### 1. Biochemistry characterization

#### Recombinant protein expression and purification

The full length coding sequence of *Erithacus rubecula* cryptochrome 4b (*Er*Cry4b) was cloned into pCold expression vector (Takara Bio, Shiga, Japan) between the restriction sites of *Spe*I and *Xho*I as described previously [34, 37] (figure S1*a*). Initially, *Er*Cry4b was expressed and purified according to the same protocol that was used for *Er*Cry4a [37]. It turned out that the protein expression protocol that worked for *Er*Cry4a did not work for *Er*Cry4b. We then optimized the protocol by co-expressing *Er*Cry4b with maltose-binding protein as the solubilizing tag. The protein expression construct is illustrated in figure S1B. The construct backbone was a gift from the Laboratory of Prof. Can Xie. To examine the protein expression level, *E. coli* cells were lyzed by ultra-sonication and subsequently centrifuged at 17,000 rpm for 40 min using the JA-25.5 centrifuge rotor (Beckman Coulter). After centrifugation, the supernatant was collected into a new tube and the cell debris pellet was dissolved in 10% Sodium Dodecyl Sulfate (SDS) detergent solution. Both supernatant and dissolved pellet samples were mixed with loading dye solution (200 mM Tris-HCl pH6.8, 1 mg/ml SDS and 50 mg/ml Bromophenol Blue, 50% glycerol and 143 mM 2-mercaptoethanol) and boiled at 95° for 5 min before loading to 12% SDS-polyacrylamide gels (figure 1C and figure S2A). Sodium Dodecyl Sulfate-Polyacrylamide Gel Electrophoresis (SDS-PAGE) were performed at 170 volts for 50-60 min. While running *Er*Cry4a and *Er*Cry4b in parallel on the same gel (figure 1D), SDS-PAGE were performed at 180 volts for 80 min.

As a further optimization, chaperones (Takara Bio) were co-expressed with *Er*Cry4b to enhance the proper folding and stability of the protein. The chaperones used in the study were groES-groEL, dnaK-dnaJ-tgrpE and tig [53, 54]. The host cells were cultured in the presence of ampicillin and chloramphenicol to the logarithmic phase. Chaperone expression was induced by adding L-arabinose to the culture at a final concentration of 13 mM. Subsequently, *Er*Cry4b protein expression was induced by 15 ^°^C cold-shock and the addition of isopropyl *β*-D-1-thiogalactopyranoside (IPTG) at a final concentration of 5 µM. Then, the protein was purified following the purification methods of *Er*Cry4a as described previously (Xu et al., 2021).

#### Antibody generation and verification

A polyclonal antibody against Cry4a was produced in rabbits (Pineda Antibody Service, Germany) using purified *Er*Cry4a protein as an antigen. To test the antibody specificity, the Cry4 rabbit antiserum was incubated with *Er*Cry1, *Er*Cry2, *Er*Cry4a, and *Er*Cry4b proteins on Western Blots. *Er*Cry1a, 1b and 2 were expressed in insect cells as previously described [55]. Cry4a and 4b were expressed in *E*.*coli* (BL21) as previously described [37]. Briefly, 10 ml insect cell culture or 30 ml cell culture was pelleted and resuspended in 1 ml Ni-binding buffer (20 mM Tris, 150 mM NaCl, 5mM Imidazole, pH 8). The cells were lysed using Dounce Homogenizers. The supernatant of cell lysates was incubated with 20 µl equilibrated Ni-NTA resin for 1 hour on a shaker platform (4 °C, 50 rpm). Subsequently, the resin was washed and eluted. The washing buffer (pH 8) contained 20 mM Tris, 150 mM NaCl and 50 mM imidazole, while the elution buffer (pH 8) contained 20 mM Tris, 150 mM NaCl and 300 mM imidazole. All buffer contained 10 mM 2-mercaptoethanol. Concentrations of eluted protein samples were measured using NanoDrop.

20 ng His-tag purified Cry proteins were loaded to SDS-PAGE gels. Meanwhile, 10 ug plain cell culture lysate was used as negative controls. The SDS-PAGE gels were run at 170 V for 45 min. Proteins on the SDS-PAGE gels were transferred to nitrocellulose membranes using Trans-Blot Turbo System (BIO-RAD). The blots were blocked using 5% milk powder in TBST (20 mM Tris, 150 mM NaCl and 0.1% Tween-20). After blocking, the blots were incubated with Cry4 rabbit antiserum (1:10,000 dilution, Pineda) or Anti-His-tag antibody (1:10,000 dilution, Invitrogen MA1-21315-HRP) at 4 °C, overnight, followed by Goat anti-Rabbit IgG secondary antibody (Invitrogen, 65-6120) incubation at room temperature for 1 hour. Blots images were obtained using SuperSignal Western Pico Plus Chemiluminescent substrate (ThermoFisher Scientific).

#### Antibody purification

*Er*Cry4a rabbit antiserum was purified using antigen-coupled CNBr-activated Sepharose 4B beads. First, *Er*Cry4a protein was coupled to CNBr resin (Cytiva 17-0430-01). 5 mg purified His-*Er*Cry4a was buffer-exchanged to coupling buffer (100 mM NaHCO_3_ and 500 mM NaCl) using PD-10 columns. 0.3 g CNBr resin were dissolved in 6 ml cold 1 mM HCl (4 °C, 15 min) and transferred to a 30-ml empty gravity-column tube to form 1 ml column bed. Subsequently, the CNBr resin was equilibrated with 5 ml coupling buffer and incubated with 5 mg His-*Er*Cry4a protein overnight at 4 °C. Next, the resin-protein mixture was washed with 6 ml coupling buffer and incubated with 5 ml 0.1 M Tris-HCl buffer (pH 8) for 2 hours at room temperature to block the uncoupled sites on the beads. Afterwards, the beads were washed three times with 2 ml acid buffer (0.1 M Acetic/Sodium Acetate, 0.5 M NaCl, pH 4) followed by 2 ml alkali buffer (0.1 M Tris-HCl, 0.5 M NaCl, pH 8.0). A final wash was performed using 10 ml PBS containing 0.02% sodium azide. The *Er*Cry4a-coupled CNBr resin (200 µl) was washed with 3 ml PBS, resuspended in 200 µl PBS and incubated with 200 µl antiserum overnight at 4 °C. Then the resin was washed with PBS buffer followed by 150 mM NaCl (pH 5) until OD280 = 0. The antibody was eluted using 150 mM NaCl (pH 2.5). The eluted antibody sample was immediately neutralized with 10% (*v*/*v*) saturated phosphate buffer. For long-term storage at -20°C, 50% (*v*/*v*) glycerol with 0.02% sodium azide was added to the antibody solution.

#### Western Blotting

For sample preparation, the protein concentration of cell lysate or immunoprecipitation elute was determined by Bradford assay. 20 to 50 µg protein samples were boiled at 95 ^°^C for 5 min in the loading buffer consisting of 200 mM Tris-HCl pH6.8, 1 mg/ml SDS and 50 mg/ml Bromophenol Blue, 50% glycerol and 143 mM 2-mercaptoethanol. Next, the protein samples were loaded on to a commercially available gradient gel (Mini-PROTEAN TGX Stain Free Gels 4-14%, Bio-Rad) and ran at 170 volts for 50-60 min. Subsequently, the proteins on the gels were transferred onto nitrocellulose membrane using a Trans-Blot Turbo apparatus (Bio-Rad) at 25 volts for 30 minutes. The transfer efficiency was examined by Ponceau staining. The membranes were then incubated with 5% milk (*v* /*v*) in TBST buffer (20 mM Tris, 150 mM NaCl, 0.1% Tween-20) at room temperature for 1 hour to block unspecific binding. After blocking, the membranes were incubated with primary antibody diluted in 2.5% (*v* /*v*) milk in TBST buffer at 4 ^°^C overnight. *Er*Cry4a monoclonal antibody from a rabbit were purified using *Er*Cry4a coupled CNBr-activated Sepharose 4B beads and diluted by 500, while *Er*Cry4a polyclonal antibody from a guinea pig [24] were diluted by 1000. On the second day, the membranes were washed 3 times with TBST and incubated with secondary antibodies in 2.5% (*v* /*v*) milk in TBST buffer at room temperature for 1 hour. Later, the membranes were washed again 3 times with TBST and rinsed with H_2_O. Last, the membranes were incubated with the chemiluminescent Western Blot reagent SuperSignal West Atto (Thermo Fisher) for 5 min in dark. Western Blot images were acquired using iBright Imaging System (Thermo Fisher Scientific).

#### Immunoprecipitation

Tissues from European robin was homogenized using glass Dounce Homogenizers in the lysis buffer of 10 mM Tris-HCl pH 7.5, 100 mM NaCl, 10% (*v* /*v*) glycerol, 1% (*v* /*v*) Triton X-100 and protease inhibitor (Roche). The tissue lysate was incubated by rotation at 4 ^°^C for 1 hour to maximum protein extraction. Protein concentration in the retina lysate was determined by Bradford assays. Immunoprecipitation was conducted according to a previous protocol [27, 36] under red light. Briefly, 20 µg purified *Er*Cry4a rabbit antibody or GFP rabbit polyclonal antibody were covalently cross-linked with 100 µl Dynabeads Protein G (Invitrogen) using 20 mM dimethyl pimelimidate dihydrochloride in 0.2 M sodium borate solution. Non-crosslinked antibodies were removed by 100 mM glycine at pH 2. Subsequently, 3 mg retina lysate was pre-cleared by incubation with empty Dynabeads Protein G beads at 4 ^°^C for 1 hour. The pre-cleared lysate was then incubated with antibody-cross-linked beads at 4 ^°^C for 1.5 hours in dark. The beads were washed to remove unspecific proteins, detergents and pH-buffering agents. 50 µl beads were eluted by boiling at 95 ^°^C and used for Western Blot as described above. The rest of the beads were saved for Mass Spectrometry measurement.

#### In-gel digestion and micro-purification

The Dynabeads-Protein G beads from the above immunoprecipitation experiments were boiled at 70 ^°^C in NuPAGE LDS sample buffer (Invitrogen) with NuPAGE sample reducing agent (Invitrogen) for 10 min. The bead samples were briefly centrifuged and supernatant from the beads samples were loaded on to Bolt 4-12% Bis-Tris gels (Invitrogen)) and ran at 160 volts for 5 min. Subsequently, the gels were stained in Quick Coomassie Stain (NeoBioTech) at room temperature for 1 to 2 hours and destained in Milli-Q ultrapure water overnight. Each lane in the gel was cut into 8 pieces from the top to the bottom according to the Coomassie blue pattern. Each gel piece was cut into small pieces (ca. 1 mm^3^). The gel pieces were washed in 80 µl 50 mM ammonium bicarbonate and then incubated in 80 µl 50 % (*v* /*v*) acetonitrile solution for 10 minutes at room temperature and 300 rpm. Subsequently, the gel pieces were de-hydrated using 100% acetonitrile. Next, the gel pieces were reduced using 10 mM dithiothreitol (30 - 45 minutes at 56 ° C) and alkylated using 55 mM iodoacetamide (30 minutes at room temperature in dark). Then the gel pieces were shrunken again using 100 % acetonitrile and incubated with 700 ng trypsin dissolved in 30 µl 50 mM ammonium bicarbonate at room temperature. 10 minutes later, excess trypsin was removed from the gel samples. The gel samples were incubated in 50 µl 50 mM ammonium bicarbonate at 37 ^°^C overnight. On the second day, the supernatant in the gel samples were collected into new tubes and acidified by adding trifluoroacetic acid to a final concentration of 0.4%. The liquid samples were further desalted using a microcolumn with the reversed-phase resin of POROS 50 R2 and OLIGO R3 packed in 10 µl pipette tips. Briefly, the samples were loaded onto the column, washed twice using 0.1 % trifluoroacetic acid and eluted using 40 µl 50% acetonitrile in 0.1 % trifluoroacetic acid and 10 µl 70% acetonitrile in 0.1 % trifluoroacetic acid. The eluate samples were dried in Speedy-vac device and re-dissolved in solvent A (0.1% *v* /*v* formic acid in H_2_O).

#### Liquid Chromatography, mass Spectrometry and data analysis

The peptide samples were analyzed using a nanoliter flow high performance liquid chromatography system (EASY-nLC, ThermoFisher Scientific) coupled to an Orbitrap Exploris 480 mass spectrometer equipped with a Nanospray Flex ion source (ThermoFisher Scientific). Specifically, the peptides samples were loaded on a two-column (Ø 75 µm) system on the EASY-nLC using 0.1% *v* /*v* formic acid (solvent A) as a mobile phase. The two-column system consisted of a 3-cm pre-column and a 18-cm analytic column, which were packed with ReproSil-Pur 120 C18-AQ resin (Dr. Maisch GmbH) of particle size 5 µm and 3 *µ*m, respectively. After sample loading, the peptides were eluted using a 60-minutes gradient. The elution gradient was initiated with a mobile phase consisting of 98% solvent A and 2% solvent B (80% acetonitrile, 0.1% formic acid). Over the first two minutes, the proportion of solvent B was gradually increased to 5%, followed by a steady increase to 25% solvent B over the next 42 minutes. Subsequently, the solvent B concentration was raised to 40% over the next 10 minutes. A rapid increase to 95% solvent B was then performed within one minute and held constant for an additional five minutes. The Exploris-480 mass spectrometer was operated in data-dependent mode at a scan range of 350 - 1600 with an orbitrap resolution of 120,000 and a normalized automatic gain control target of 200%. A tandem mass spectrometry (MS/MS) technique was used to break down peptide ions into smaller fragments. The data-dependent MS/MS scan was performed at an isolation window of 1.2 m/*z* and collision energies of 35% with a maximum injection time of 100 ms. To identify the peptides, all hereby created MS/MS spectra were loaded into Proteome Discoverer (version 2.5.0.400, Thermo Fisher Scientific). Peptide search was performed in the proteomic database of *Erithacus rubecula* proteome (UniProt UP000529965) and a FASTA file of *Er*Cry4b sequence using Mascot search engine. The search parameters were ±0.1 Da fragment mass tolerance and ±8 ppm for precursor mass tolerance. The maximum number of missed cleavages was set to 2. Carbamidomethyl on cysteine was set as a static modification and oxidation on methionine was set as a variable modification. The minimum peptide length was set to 4 amino acid residues. The result was filtered to 1% false discovery rate on protein level using the Percolator algorithm integrated in Thermo Proteome Discoverer (Version 2.5.0.400).

## 2 Molecular dynamics simulations

### Simulation setup

The initial protein structures for both *Er*Cry4a and *Er*Cry4b were determined using the unrelaxed predictions from the OpenFold [56] implementation of the AlphaFold [57]. Four template structures were selected by the algorithm to refine each model. For *Er*Cry4a, the algorithm identified two crystal structures from pigeon cryptochrome (pdb IDs: 6PTZ and 6PU0) and two crystal structures from drosophila cryptochrome (pdb IDs: 4JZY and 4GU5) as suitable templates. The *Er*Cry4b modelling procedure also made use of the pigeon cryptochrome structures (pdb IDs: 6PTZ and 6PU0) and complemented it with structures from green alga cryptochrome (pdb IDs: 6FN0 and 5ZM0). When included, the non-covalently bound cofactor FAD was positioned by aligning the equilibrated structure of *Er*Cry4a from an earlier study [58] with the respective model using the STAMP algorithm [59] as implemented in the MultiSeq tool [60] of VMD [61].

Protonation states for ionizable residues were determined using PROPKA 3 [62, 63] and neutral histidine were assumed to be in their D-histidine isomer form. The cryptochromes were prepared to represent the so-called dark state which means that the flavin cofactor was expected not to be light activated, i.e. with an electrostatically neutral flavin part in the singlet state and no hydrogen at the N5 atom. Consequently, the surrounding tryptophan residues, which have been found to donate electrons to the light-activated flavin [39], were electrostatically neutral as well. In all simulation, the proteins were solvated in a cubic water box of explicit water molecules with a side length of 130 Å to ensure that the flexible C-terminal regional is allowed to fluctuate freely without causing boundary artifacts. After pressure equilibration, the waterbox shrunk to a cube with side length of 127.5 Å. The system was neutralized and ionized using NaCl ions to match a salt concentration of 0.15 mol/L. System preparation was executed in VMD [61].

Molecular dynamics (MD) parameters for the protein and solvent were defined by the CHARMM36m force field which includes CMAP corrections [64, 65, 66] in combination with the TIP3P water model which was used for the derivation of the parameter set. Parameters for the FAD were adopted from an earlier study [67]. The simulations were run using NAMD 3.0b4 [68]. Non-bonded interactions were linearly switched off between 10 - 12 Å and long-range electrostatics were treated via a smooth Particle-Mesh-Ewald [69] algorithm employing a grid spacing of 1 Å. Non-bonded interactions were not applied to atoms within the molecule that were separated by less than four bonds. Non-bonded interactions were computed at every step, but long-electrostatics were only computed every second step. A Langevin thermostat allowed for temperature control at 300 K with a damping constant of 5.0 ps^-1^ acting on all non-hydrogen atoms. Pressure control, where active, made use of the Nose-Hoover Langevin piston algorithm [70, 71] with a target pressure of 1.01325 bar, a period time of 200 fs and a decay time of 50 fs.

Equilibrium MD simulations started with conjugate gradient minimization for 20,000 steps followed by 5 ns of simulation employing a 1 fs timestep in the NPT ensemble and isotropic cell size fluctuations. Afterwards, all bonds involving hydrogen atoms were fixed using the SETTLE [72] algorithm for water molecules and the SHAKE algorithm [73] for all other hydrogens, which allowed to simulate using a timestep of 2 fs. With these settings, the simulations were continued for another 5 ns in the NPT ensemble and finally 500 ns in the NVT ensemble. The structures for Cry4a and Cry4b were both simulated with or without a bound FAD molecule and six replicas for each of the four simulations. The workflow was automatized with the NAMD AutoConf wrapper program.

The analysis was carried out using tools from VMD [61] and the MDAnalysis python library [74, 75]. General large language models assisted in data curation and analysis, but no code was run without thorough code review of a language-proficient developer.

### Definitions

Sequence inclusion: K178-R206 in the *Er*Cry4b sequence. Purely sequence based alignment of *Er*Cry4a vs *Er*Cry4b results in a continuous series of gaps in the aligned *Er*Cry4a sequence. Apart from the inclusion, the two sequences only differ in one position of the alignment which corresponds to L289 in Cry4a and W318 in *Er*Cry4b.

Structural inclusion: E179-D214 in the *Er*Cry4b sequence. Structural sequence alignment Russell and Barton, 1992 of *Er*Cry4a and *Er*Cry4b results in a region that is dominated by many gaps in the aligned *Er*Cry4a sequence. The first of these gaps relates to E179 in the *Er*Cry4b sequence and the last gap corresponds to D214 in the *Er*Cry4b sequence.

PBL: phosphate binding loop; G227-P244 in Cry4a; G256-P273 in *Er*Cry4b. This region was defined as the sequence that matches the part of the loop that was unresolved in the pigeon cryptochrome crystal structure by Zoltowski *et. al*. (pdb ID: 6PTZ). Additionally, the structure prediction shows reduced certainty for this sequence. The amino acid sequence of the PBL is identical in *Er*Cry4a and *Er*Cry4b.

C-terminal: R497-E527 in *Er*Cry4a; R526-E556 in *Er*Cry4b. The last 30 residues are unresolved in the pigeon crystal structure (pdb ID: 6PTZ) and the structure prediction portrays very high uncertainty.

### Principal Component Analysis (PCA)

PCA allows to find a projection of the initial coordinate system on to a new coordinate system in which the eigenvalues can be used to sort the new coordinate axis in the order of their importance with regards to describing the trajectories that are being analyzed. In this case, it was also aimed that the PCA of the four different simulation setups (*Er*Cry4 + FAD, *Er*Cry4a, *Er*Cry4b + FAD, *Er*Cry4b) are comparable and that the differences in motion are limited to those that are expected to be relevant for the stability of the protein and its ability to bind FAD. Therefore, the first step of the PCA consisted of reducing the analyzed trajectories to the non-hydrogen atoms of the protein excluding the structural inclusion, the PBL and the C-terminal. The excluded regions are expected to undergo major motions and possible conformational changes. However, due to a physically large distance towards the FAD binding pocket neither the structural inclusion nor the C-terminal itself necessarily have an impact on the ability of the protein to bind FAD. The question to be addressed is instead, whether the possibly different motions of the structural inclusion or the C-terminal could induce other conformational changes that do affect the expected binding properties. Residue L289 in *Er*Cry4a and residue W318 in *Er*Cry4b needed special consideration, since this single-point mutation is an additional (and the only) difference between the protein apart from the structural inclusion. Only the heavy atoms of the backbone as well as the C-*β*, C-*γ* and the two C-*δ* atoms were included from those residues. Prior to PCA calculation, all trajectories have been aligned with respect to the backbone atoms of the included regions.

### Solvent accessibility of the FAD binding pocket

Residues were selected by starting with the energy minimized structures of the first replica of the simulation that contained FAD and then including all amino acid residues for which at least one atom was within (3.0, 3.5 4.0, 4.5, 5.0) Å of the FAD. By choosing a range of radii for the analysis, the result is less dependent on slight structural differences and informs on the spread of values. The solvent accessible surface area (SASA) was measured in VMD using a probe radius of 1.4 Å. Since the number of residues in the shells varies and since different amino acids possess different maximal SASA, the SASA was normalized using the *Theoretical ALL* result for the maximal SASA of the amino acids as computed by Tien et al., 2013, who also employed a probe radius of 1.4 Å. The normalized SASA is then interpreted as the relative solvent exposure. A sliding window averaging over *±*25 ns was performed on every dataset and from the averaged data, the median with respect to the different solvent shell radii was highlighted as a thick line in figure 2.

### Transcript analysis

European robin tissue was collected in September 2019. Tissue mRNA was extracted using TRIzol (Invitrogen, Cat. No. 15596-026) following instructions of the manufacturer. Total mRNA was cleaned up using LiCl. The mRNA samples were sent to Cold Spring Harbor laboratories (NY, USA) for isoseq on a Pacbio sequencer. A high quality ISOSeq transcriptome of European robins was generated for 8 tissue samples (retina, muscle, brain, lung, liver, heart, skin and testis). The sequencing of full transcripts allowed us to investigate the different isoforms available in the transcriptome and compare their absolute numbers. The sequences were barcoded with the IsoSeq barcodes v1 and sequenced on a PacBIO Sequel II. PacBIO CCS (4.0.0), lima (1.10.0) and refine (3.4.0) where used for consensus read generation, adapter removal, barcode identification and PolyA trimming. Afterwards, pbmm2 (1.19.2) was used to align the full-length transcripts to the reference genome (GeneBank ID GCA 903797595.1). Sequences were traced back to their tissue of origin using the barcodes.The location of *ErCRY4* was identified using BLAST. Transcripts were investigated visually using IGV [76]. Any transcripts that did not cover the full coding sequence of *Er*Cry4a were discarded. The remaining transcripts were assigned to either *ErCRY4a, ErCRY4b* or one of the newly identified spliced variants of *ErCRY4*. For each splice variant and tissue, the number of transcripts was counted and plotted using a custom python script.

## Author contribution

J.X. conceived the project and designed the experiments. J.X. purified the antibody. J.X. and A.B.P.A. executed recombinant protein expression and bird tissue co-immunoprecipitation experiments. J.X. conducted mass spectrometry measurement and data analysis. J.H. conducted computational simulations. G.L. and A.N. worked on the robin transcriptome. T.R. and O.N.J. provided crucial support in mass spectrometry measurement and data analysis. J.S. assisted in protein expression. B.S. and L.S. collected the bird tissue. K.D. helped on antibody generation. R.B. and T.K. advised on protein expression, M.L., I.A.S., and H.M. contributed constructive discussion. J.X., J.H., and G.L. plotted the figures and wrote the initial version of the manuscript. All authors made significant contributions to the final version.

## Ethics statement

Bird catching was performed based on a permit from the Lower Saxony State Department for Waterway, Coastal and Nature Conservation (D7.2220/18).

## Acknowledgements

The authors would like to thank the Lundbeck Foundation (Postdoc fellowship grant number: R402-2022-1372, awarded to J.X.); Novo Nordisk Foundation (INTEGRA, grant number NNF20OC0061575 to O.N.J.); European Research Council (under the European Union’s Horizon 2020 research and innovation programme, grant agreement no. 810002, Synergy Grant: ‘QuantumBirds’, awarded to H.M.); Volkswagen Foundation (Lichtenberg Professorship to I.A.S.), the Deutsche Forschungsgemeinschaft (SFB 1372: Magnetoreception and Navigation in Vertebrates, no. 395940726 to H.M., M.L., I.A.S., K.D. and A.N.); TRR386/1-2023 Hyperpolarization in molecular systems HYP*MOL, no 514664767 to I.A.S.), the Ministry for Science and Culture of Lower Saxony (Simulations Meet Experiments on the Nanoscale: Opening up the Quantum World to Artificial Intelligence (SMART) and Dynamik auf der Nanoskala: Von kohärenten Elementarprozessen zur Funktionalität (DyNano)). Computational resources for the simulations were provided by the CARL Cluster at the Carl-von-Ossietzky University Oldenburg, which is supported by the DFG and the ministry for science and culture of Lower Saxony. The authors gratefully acknowledge the computing time made available to them on the high-performance computers HLRN-IV at GWDG at the NHR Centers NHR@Göttingen. These Centers are jointly supported by the Federal Ministry of Education and Research and the state governments participating in the NHR. J.X. thanks David Keays, Alexandra Vilceanu and Spencer D.Balay for advice on immunoprecipitation experimental protocols. We are grateful to Vibeke Jørgensen, Lene Jakobsen, and Elisa Le Boiteux for help on mass spectrometry measurements.

