## Supplementary Information for "Cryptochrome 4b protein is likely irrelevant for the radical pair based magnetoreception in the European robin"

### Supplementary information for: Cryptochrome 4b protein is unlikely relevant to the radical pair based magnetoreception in the European robin

Jingjing Xu[1#], Alisha Bhanu Pattani Ameerjan[2], Jonathan Hungerland[3], Georg Langebrake[3, 4], Tina Ravnsborg[1], Ole N. Jensen[1], Jessica Schmidt[2], Rabea Bartölke[2], Takaoki Kasahara[2], Baladev Satish[2], Leonard Schwigon[2], Karin Dedek[2], Arne Nolte[2], Miriam Liedvogel[2, 4], Ilia A. Solov'yov[3,5,6], and Henrik Mouritsen[2,6]

[1]Department of Biochemistry and Molecular Biology, University of Southern Denmark, Campusvej 55, 5230 Odense, Denmark

[2]Institute of Biology and Environmental Sciences, University of Oldenburg, Carl-von-Ossietzky-Str. 9-11, 26129 Oldenburg, Germany

[3]Institute of Physics, University of Oldenburg, Carl-von-Ossietzky-Str. 9-11, 26129 Oldenburg, Germany

[4]Institute of Avian Research, An der Vogelwarte 21, 26386 Wilhelmshaven, Germany

[5]Center for Nanoscale Dynamics (CENAD), University of Oldenburg, Ammerländer Heerstr. 114-118, 26129 Oldenburg, Germany

[6]Research Centre for Neurosensory Science, University of Oldenburg, Carl-von-Ossietzky-Str. 9-11, 26129 Oldenburg, Germany

### 1 Biochemistry

**Establishing an efficient protein expression system for *Er*Cry4b**

The first step towards biochemical characterization is to produce large quantities of *Er*Cry4b protein. A pilot protein expression was conducted using the protocol previously established for *Er*Cry4a [1]. However, *Er*Cry4b was poorly expressed using the same protocol as that for *Er*Cry4a (figure S1*a* and figure S1*d*). Due to the low protein expression level, *Er*Cry4b was only detectable by Western blotting (figure S1*d*), a signal amplification technique [2]. The result showed that very little *Er*Cry4b was soluble (figure S1*d*, Lane S), and majority of *Er*Cry4b were present in the insoluble pellet fraction (figure S1*d*, Lane P). To enhance the protein solubility, a solubilizing tag of maltose binding protein (MBP) was introduced into the protein expression construct (figure S1*b*). But this modification only slightly improved the protein expression (figure S1*e*). Further optimization was attempted by co-expressing *Er*Cry4b with various chaperones. Among different chaperones, GroES-GroEL chaperone greatly improved the protein expression, as observed by the protein band in Lane S in figure S1*f*.

**Antibody verification**

To verify the new antibody generated in the present study, we conducted Western blotting using the antiserum against different Cry proteins from robins. The results showed that the new *Er*Cry4 antibody recognized both *Er*Cry4a and *Er*Cry4b as expected. Moreover, the antibody specificity is high, as it did not bind to *Er*Cry1 or *Er*Cry2 proteins.

**Purified *Er*Cry4b protein as a positive control in mass spectrometry**

To establish an efficient bottom-up mass spectrometry methodology for the present study, purified *Er*Cry4b protein was used as a positive control. Upon fine tuning on instrument parameters, a high confidence peptide spectra match was achieved for the unique and specific peptide of *Er*Cry4b VKDVLCLALHEER. Table S1 shows that the measured $a^+$, $b^+$ and $y^+$ ions match to the theoretical mass. The high confidence and good match suggest that the MS methodology is powerful to detect the *Er*Cry4b-specific peptide. Thus, the established MS method was applied to robin retina immunoprecipitation samples. *Er*Cry4b unique peptide was missing in retina immunoprecipitation samples (figure S4). All the identified 12 peptides belong to *Er*Cry4a protein. figure S5 indicates that the *Er*Cry4b specific peptide was not found in cerebellum or liver.

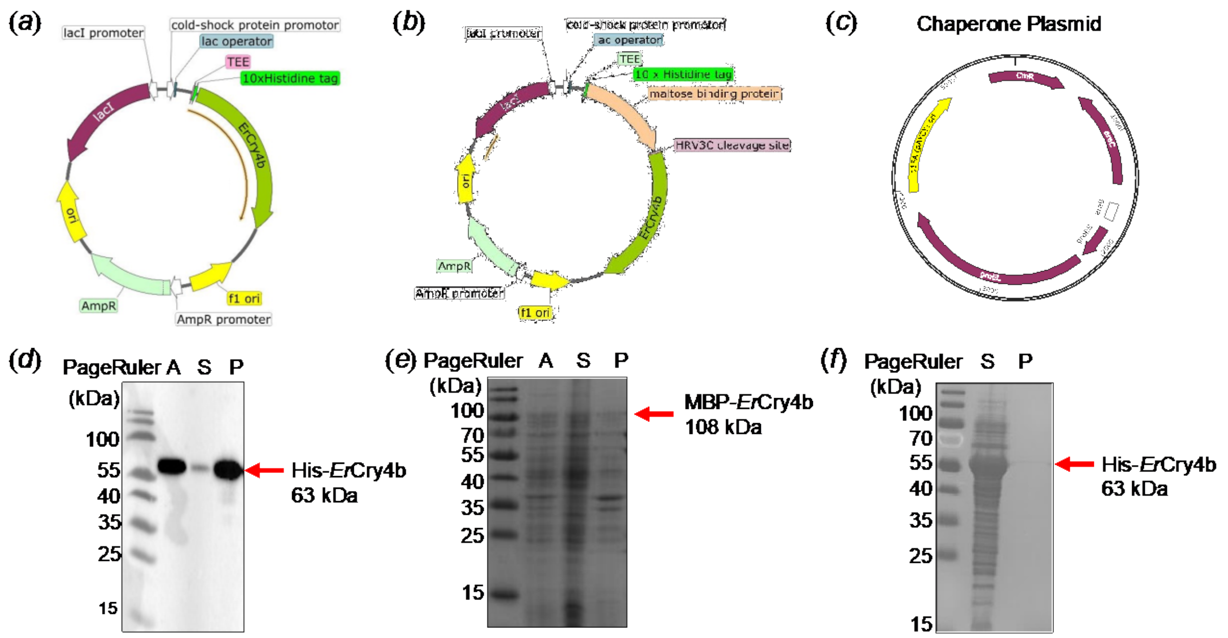


**Figure S1. Establishing an efficient protein expression system for *Er*Cry4b.** (*a* - *c*) Graphic illustration of *Er*Cry4b protein expression and chaperone plasmids used in the present study. Key genetic elements are marked in different colors. AmpR stands for Ampicillin resistance. The element of ‘f1 ori’ stands for f1 bacteriophage origin of replication. lacI encodes Lac repressor protein, which is used to regulate protein expression. ’TEE’ stands for a translation enhancing element consisting of the amino acid residues MNHKV. (*d* - *f*) Protein expression test using different plasmids above. Lane A: *E. coli* cells after induction; Lane S: soluble fraction (supernatant); Lane P: insoluble fraction (pellet). The expected molecular weight of His-*Er*Cry4b and MBP-*Er*Cry4b are 63 kDa and 108 kDa, as indicated by red arrows. Note, D is a Western Blot images, while E and F are Coomassie blue staining images of sodium dodecyl sulfate–polyacrylamide gels.

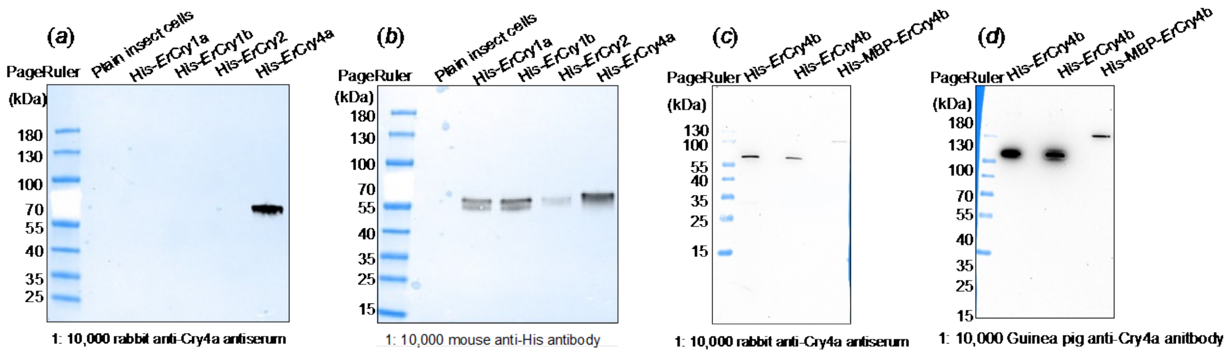


**Figure S2. Antibody specificity test against the different cryptochromes found in robin analyzed by Western Blotting (WB).** (*a*, *c*) Blots using *Er*Cry4 rabbit antiserum generated in the present study show that the newly developed *Er*Cry4 antibody recognize both *Er*Cry4a and *Er*Cry4b, but not *Er*Cry1a, *Er*Cry1b or *Er*Cry2. (*b*, *d*): WB using His-tag antibody or previously reported Guinea pig Cry4 [3] antibody as a positive control to confirm protein expression.

**Figure S3. A single Cry4 protein in robin retina.** GFP antibody was used as a negative control to verify specificity.

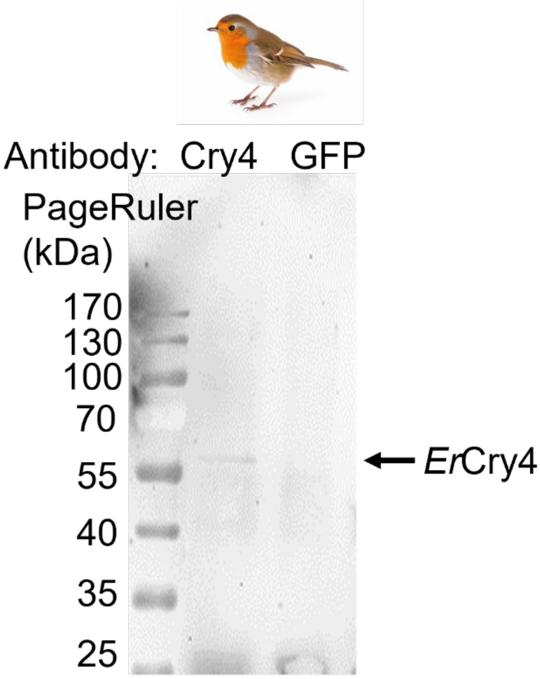

**Table S1. Ion series of *Er* Cry4b-specfic peptide fragment matches.** The brown and green colors mark the ions that were detected, i.e., the measured mass matches to the theoretical mass.

| Sequence | $a^+$ ions | $a^{2+}$ ions | $b^+$ ions | $b^{2+}$ ions | $y^+$ ions | $y^{2+}$ ions |
|---|---|---|---|---|---|---|
| V | 72.08078 | 36.54403 | 100.07569 | 50.54148 | | |
| K | 200.17574 | 100.59151 | 228.17065 | 114.58897 | 1482.77332 | 741.89030 |
| D | 315.20268 | 158.10498 | 343.19760 | 172.10244 | 1354.67836 | 677.84282 |
| V | 414.27110 | 207.63919 | 442.26601 | 221.63664 | 1239.65142 | 620.32935 |
| L | 527.35516 | 264.18122 | 555.35007 | 278.17868 | 1140.58300 | 570.79514 |
| C-Carbamidomethyl | 687.38581 | 344.19654 | 715.38072 | 358.19400 | 1027.49894 | 514.25311 |
| L | 800.46987 | 400.73857 | 828.46479 | 414.73603 | 867.46829 | 434.23778 |
| A | 871.50699 | 436.25713 | 899.50190 | 450.25459 | 754.38423 | 377.69575 |
| L | 984.59105 | 492.79916 | 1012.58596 | 506.79662 | 683.34711 | 342.17720 |
| H | 1121.64996 | 561.32862 | 1149.64488 | 575.32608 | 570.26305 | 285.63516 |
| E | 1250.69255 | 625.84992 | 1278.68747 | 639.84737 | 433.20414 | 217.10571 |
| E | 1379.73515 | 690.37121 | 1407.73006 | 704.36867 | 304.16155 | 152.58441 |
| R | | | | | 175.11895 | 88.06311 |

**Figure S4. Overview of *Er*Cry4 peptides identified in the robin retina immunoprecipitation sample**. Identified peptides were highlighted in yellow shades. The *Er*Cry4b specific peptide was indicated in the red dash line box.

MLHRTIHLFRKELRLHDNPVLLAALQSSEALYPVYILDRAFLTSSMHIGALRWHFLLQ
SLEDLHKNLCQLGSCLLVIQGEYETVLRDHIQKWSITQVTLDAEMEPFYKEMEANIQ
CLGAELGFEVLSLGSHSLYDTQRILDINGGSPPLTYKRFLHILSLLGDPEVPVRPNLT
AEDFQKEGLSSCLPGLEKKLRVKDVLCLALHEERRCSAPDPDLAECYRVPLPVDLK
ISPENLSPWRGGETEGLQRLEQHLTDQGWVASFTKPRTIPNSLLPSTTGLSPYFSM
GCLSVRTFFYRLSNIYAQAKHHSLPPVSLQGQLLWREFFYTVASATPNFTQMAGNP
ICLQICWYKDAERLHKWKMAQTGFPWIDAIMTQLRQEGWIHHLARHAVACFLTRG
DLWISWEEGMKVFEELLLDADYSINAGNWMWLSASAFFHQYTRIFCPVRFGKRTD
PQGNYIRKYLPILKNFPSKYIYEPWTASEEEQKQAGCIIGRDYPFPMVNHKEASDH
NLQLMRQVREEQHRTAQLTRDDADDPMEIKVKRDHTEENISKGKVARTTE*

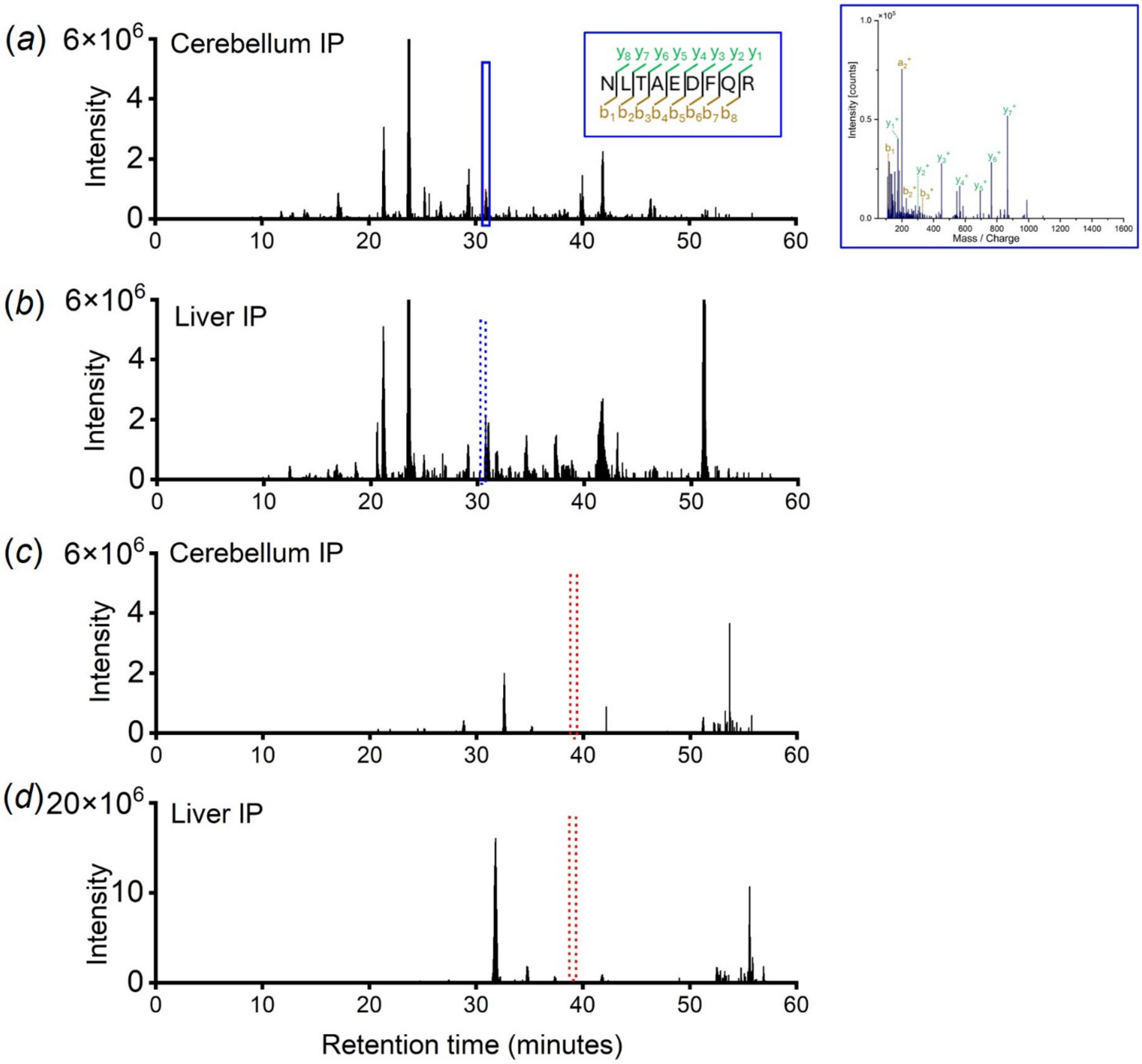


**Figure S5. Detecting *Er*Cry4b in robin cerebellum and liver.** *Er*Cry4a and 4b common peptide was found in cerebellum (*a*), but not in liver (*b*). *Er*Cry4b specific peptide was not found either in cerebellum (*c*) or liver (*d*).

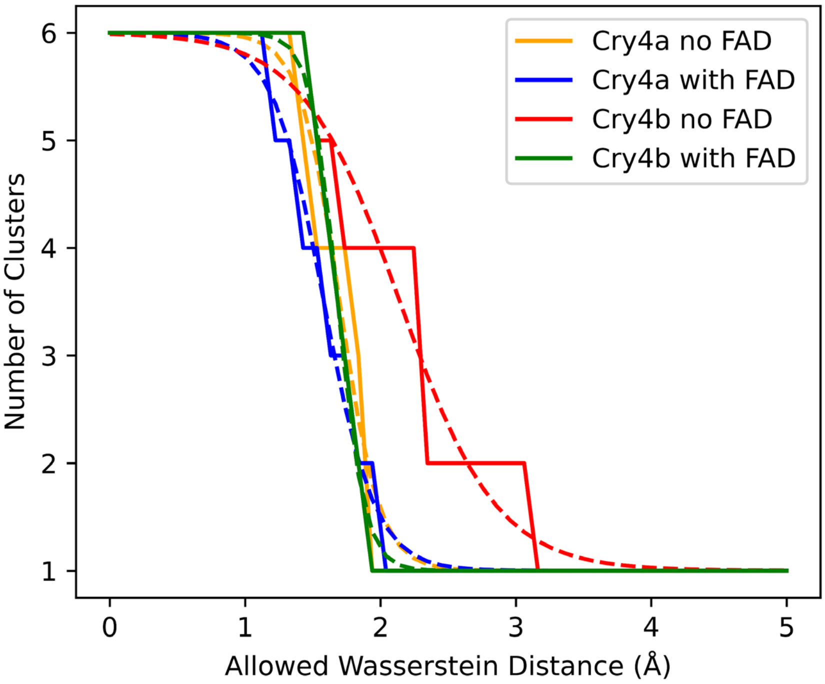


**Figure S6. Clustering analysis based on the wasserstein analysis together with fits according to Eq. (S1).**

#### 2 Molecular Dynamics simulations

##### Wasserstein Clustering Analysis

The PCA results suggest that the *Er*Cry4a trajectories are more similar, independent of the presence of FAD, while there are differences in the *Er*Cry4b trajectories. A quantification of this observation was executed by means of a clustering analysis of the Wasserstein distance. First, each trajectory was projected on to the first 100 principal components. Then, the wasserstein distance between every possible pair of two projected trajectories was computed using the SciPy python library [4], leading to one number for the distance between the trajectories. For the sake of comparison with other trajectories and distance measurements, this number was divided by the total number of atoms resulting in units of $Å^2$/atom. The different sets of trajectories were combined into groups in which a clustering analysis was performed. The number of clusters N within a group of L trajectories, was counted for different maximally allowed wasserstein distances of $d$ $Å^2$/atom. While for $d \to 0$, every trajectory in the group is alone in its cluster and $N = L$, the case $d \to \infty$ would sort all trajectories into the same cluster and $N = 1$. For the sake of condensing the results, a sigmoid-like function was fitted to the evolution of $N$ with $d$ which is

$$N(d) = L - \frac{L-1}{1+\exp(-k(d-d_0))} \tag{S1}$$

Finally, the results may be summarized in form of the Table S2. The values of $d_0$ provide a measurement of the distance of a set of trajectories within the group. Errors were estimated as the maximal deviation in the parameters that can be achieved by removing any single trajectory from the group. By grouping all six replicas of *Er*Cry4a with FAD, their distance is 1.58 Å/atom and grouping the six replicas of *Er*Cry4a without FAD results in 1.70 Å/atom. Forming a larger group of all simulations of *Er*Cry4a into a total of 12 trajectories reduces the distance to 1.51 Å/atom. Since the distance within the combined group is not larger than the distance in the individual groups, this lets one conclude that the presence of FAD does not change the covered phase space of the *Er*Cry4a protein. For *Er*Cry4b, different observations are made. Firstly, the distances are larger overall, indicating that *Er*Cry4b covers a larger portion of phase space, which leads to greater distances between replicas. In other words, *Er*Cry4b, even in the PCA space based on coordinate intersection, is more flexible and displays greater movements. Secondly, the combined group of all *Er*Cry4b trajectories has an increased distance with respect to the FAD containing *Er*Cry4b group. It suggests that the removal of FAD leads to conformational changes within the protein that would not happen in the presence of FAD. While *Er*Cry4b adapts its conformation to the presence of FAD, *Er*Cry4a retains the same phase space independent of FAD, making it more likely for FAD to bind in *Er*Cry4a once the two molecules encounter each other.

**Table S2. Fit parameters for the sigmoid functions according to the equation S1**. Errors are estimated by computing the maximal deviation that can be achieved by removing a single trajectory from the analysis. Units are $Å^2$ /atom. Note, that the distance matrix is symmetric, which makes values below the diagonal redundant.

| **$d_0$** | Cry4a with FAD | Cry4a no FAD | Cry4b with FAD | Cry4b no FAD |
|---|---|---|---|---|
| Cry4a with FAD | 1.58 ± 0.15 | 1.51 ± 0.04 | 1.68 ± 0.05 | 1.86 ± 0.11 |
| Cry4a no FAD | | 1.70 ± 0.17 | 1.73 ± 0.08 | 1.91 ± 0.10 |
| Cry4b with FAD | | | 1.68 ± 0.08 | 1.78 ± 0.07 |
| Cry4b no FAD | | | | 2.14 ± 0.25 |
| **k** | | | | |
| Cry4a with FAD | 5.27 ± 0.82 | 7.48 ± 1.24 | 6.09 ± 0.92 | 3.33 ± 0.67 |
| Cry4a no FAD | | 6.83 ± 2.50 | 6.91 ± 1.36 | 3.68 ± 0.80 |
| Cry4b with FAD | | | 10.00 ± 0.66 | 5.97 ± 1.98 |
| Cry4b no FAD | | | | 2.79 ± 1.08 |

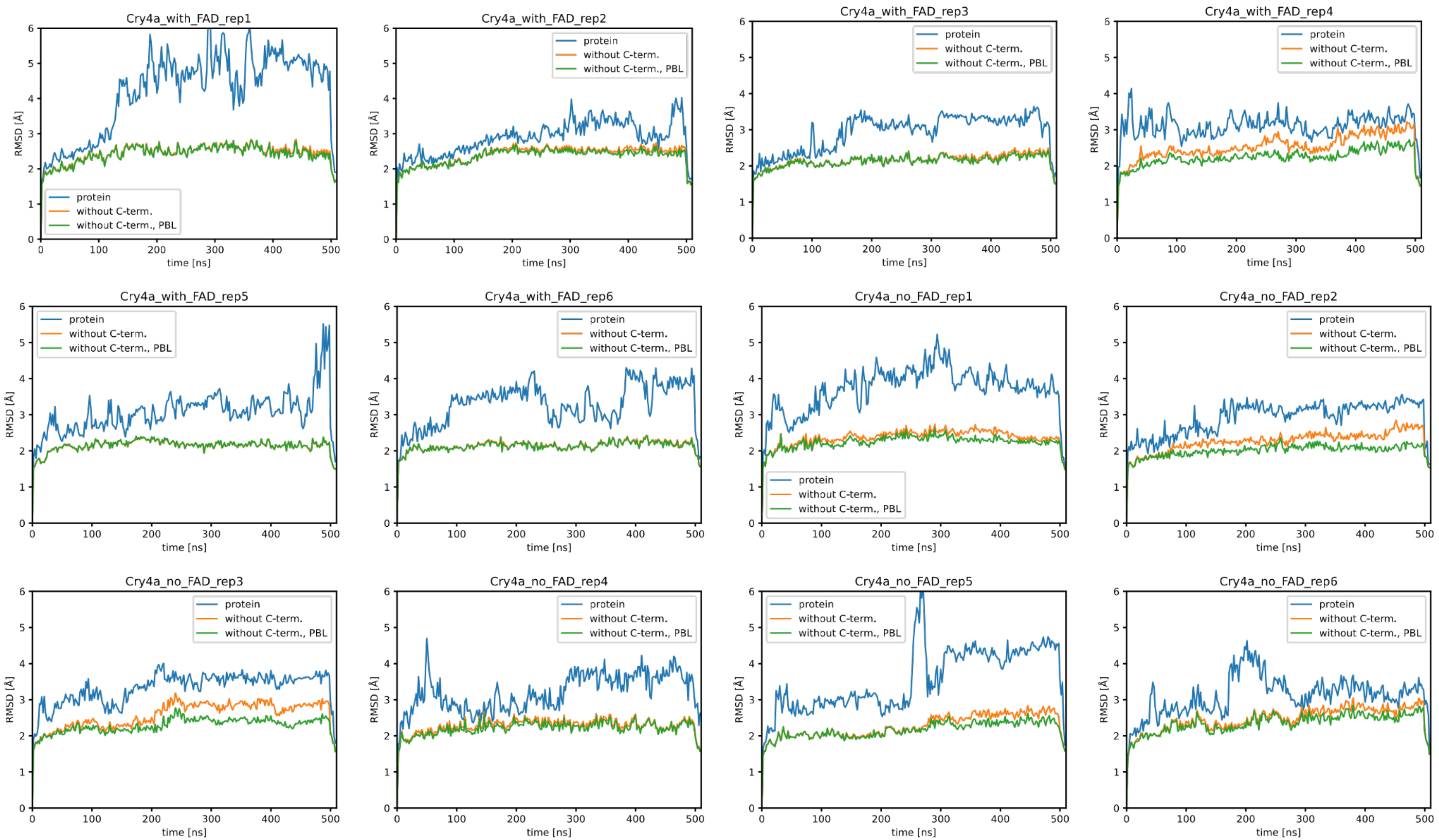


**Figure S7. Root mean square deviation (RMSD) over time of *Er*Cry4a protein with or without FAD.** Prior to RMSD calculation, each protein structure was aligned with respect to the structure after minimization using the backbone atoms without C-terminal, PBL and structural inclusion. Each RMSD analysis was repeated 6 times.

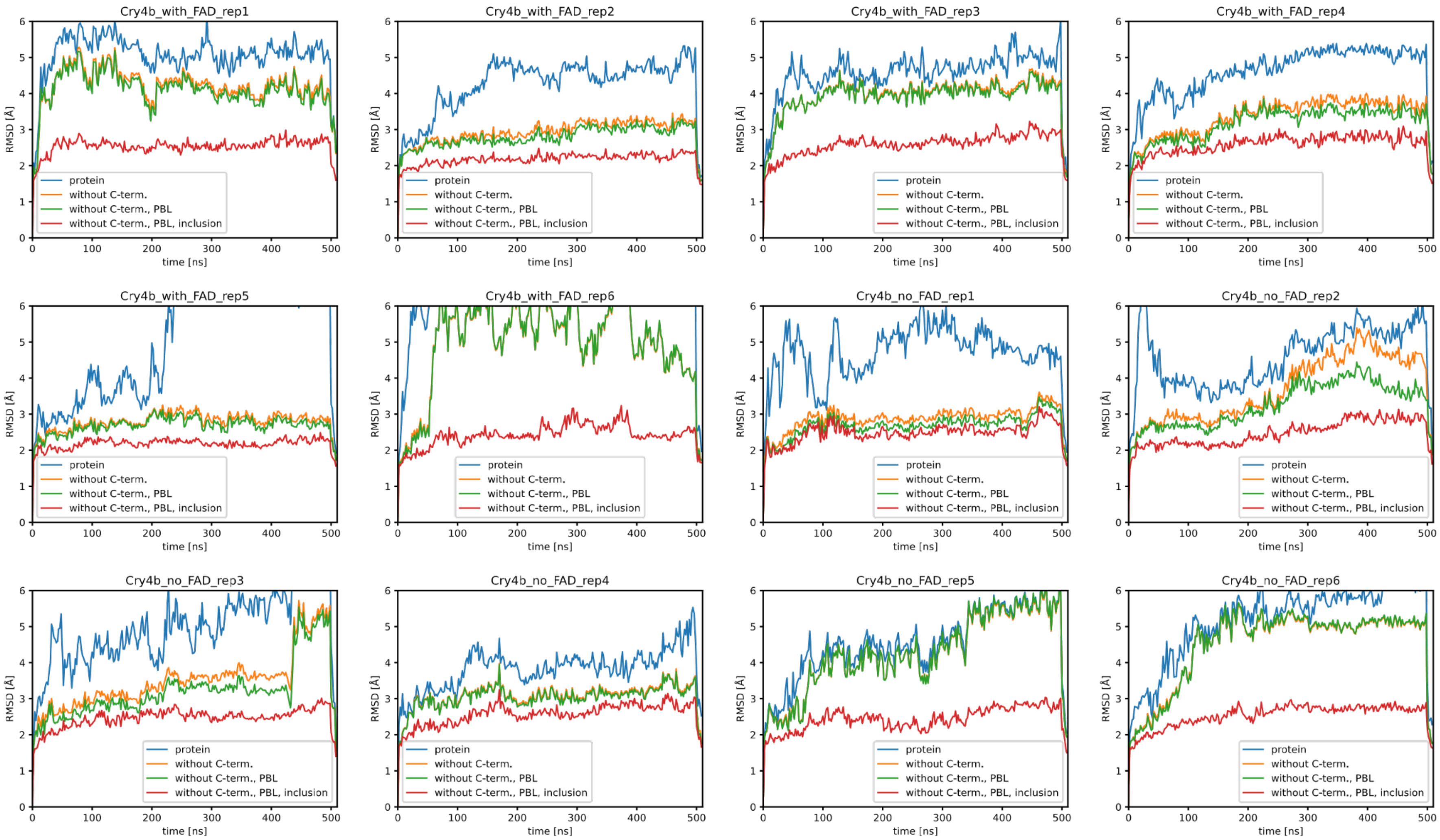


**Figure S8. Root mean square deviation (RMSD) over time of *Er*Cry4b protein with or without FAD.** Prior to RMSD calculation, each protein structure was aligned with respect to the structure after minimization using the backbone atoms without C-terminal, PBL and structural inclusion. Each RMSD analysis was repeated 6 times.

#### 3 Transcriptomics

Next to the tissues investigated in the main text, we have also sequenced transcriptomes of muscle, lung, heart and testis. We also find additional transcript types with additional introns, but which only occur a single time in all tissues. These transcripts are characterized by the following features:

n7: Intron between *ErCRY4a* exons 3 and 4.

n8: Like n5 but with an additional intron between *ErCRY4a* exons 1 and 2.

n9: Introns between *ErCRY4a* exons 2 and 4, as well as between 9 and 10.

n10: Like n5 but with a 227 bp and a 15bp insertion in *ErCRY4a* exon 7.

n11: Like n2 but with an additional exon between *ErCRY4a* exons 9 and 10.

n12: Like n4 but with a longer b exon.

The full account of all transcript types in different organs were shown in Table S3.

**Table S3. Number of different Cry4 transcripts found in various organs**

| Transcript type | Retina | Cerebellum | Liver | Skin | Muscle | Lung | Heart | Testis |
|---|---|---|---|---|---|---|---|---|
| a | 12 | 5 | - | 7 | 1 | 6 | 2 | 3 |
| b | 1 | 3 | 1 | 4 | - | 3 | 2 | 3 |
| n1 | - | - | - | 6 | - | - | - | - |
| n2 | 5 | 2 | - | 36 | - | 2 | 1 | - |
| n3 | 5 | 4 | - | 26 | - | 1 | 1 | 1 |
| n4 | - | - | - | 1 | - | - | 1 | 1 |
| n5 | - | - | - | 4 | - | 1 | - | - |
| n6 | - | 1 | - | 1 | - | - | - | - |
| n6 | - | 1 | - | - | - | - | - | - |
| n7 | - | - | - | 1 | - | - | - | - |
| n8 | - | - | - | 1 | - | - | - | - |
| n9 | - | - | - | 1 | - | - | - | - |
| n10 | - | 1 | - | - | - | - | - | - |
| n11 | - | - | - | 1 | - | - | - | - |
| n12 | - | - | - | 1 | - | - | - | - |
